# Hormonal contraceptive intake during adolescence and cortical brain measures in the ABCD Study

**DOI:** 10.1101/2024.11.07.622544

**Authors:** Carina Heller, Elvisha Dhamala, Katherine L. Bottenhorn, Megan M. Herting, Brian Bossé, Jennifer S. De La Rosa, Leslie V. Farland, Alicia M. Allen, Claudia Barth, Nicole Petersen

**Author notes:** <u>Corresponding author:</u> Nicole Petersen, Department of Psychiatry and Biobehavioral Sciences, David Geffen School of Medicine at UCLA, Los Angeles, USA. <u>Author contributions:</u> CH was responsible for methodology, visualization, and writing – original draft. ED contributed to formal analysis and writing – review & editing. KB contributed to formal analysis and writing – review & editing. MMH contributed to writing – review & editing. BB contributed to visualization and writing – review & editing. JSDLR contributed to writing – review & editing. LVF contributed to writing – review & editing. AMA contributed to writing – review & editing. CB contributed to writing – review & editing. NP was responsible for conceptualization, methodology, formal analysis, visualization, resources, supervision, writing – review & editing. <u>Competing interest:</u> The authors declare no financial or non-financial competing interest. <u>Data availability:</u> The dataset that supports the findings of this analysis is under controlled access, available on the ABCD Data Repository through the NIMH Data Archive (NDA). (https://abcdstudy.org). <u>Code availability:</u> All analysis scripts are publicly available on our GitHub repository: https://github.com/HumanBrainZappingatUCLA/ABCDOCPS/. **Brief Communication** A Brief Communication is a format intended for reporting of timely new results that, while limited in scope, are of substantial clinical or public health importance, and that therefore need to be quickly vetted and shared.

## Abstract

Adolescence is a critical period in human development, marked by rapid changes in brain structure and function, influenced, in part, by sex steroids. During adolescence, females may initiate the use of hormonal contraception, which suppresses ovarian production of estradiol and progesterone. Here, we examined the impact of hormonal contraceptive use on adolescent brain structures using magnetic resonance imaging data from 1,234 individuals from the ABCD Study (average age = 14 years). We investigated differences in cortical thickness, surface area, and volume in hormonal contraceptive-users compared to non-users. Statistically significant differences were observed in several regions, and differences in cortical thickness of the paracentral gyrus survived familywise error correction. Estradiol, testosterone and DHEA levels negatively correlated with cortical thickness, surface area, and volume across both hormonal contraceptive-users and non-users only at an uncorrected threshold. These findings highlight the need to further study longitudinal effects of hormonal contraceptives and endogenous hormone changes on brain development in the ABCD Study and related datasets as these data become available.

## Main

Adolescence is a critical period in human development, marked by a dynamic cascade of changes in both brain structure and function^1^. During this transformative time, neuromaturational processes such as myelination and synaptic pruning actively shape the brain^2^, with maturation occurring in a back-to-front pattern^3^, and protracted development of prefrontal structures by the mid to late 20s^4^. Pubertal increases in sex steroids, including estrogens, progesterone, and testosterone, play a key role in driving neurodevelopmental trajectories crucial for cognitive and emotional maturation^5^. Previous research has linked fluctuations in these hormones to variations in gray and white matter during adolescence^6^, with notable sex differences^7^. These hormonal shifts are also thought to potentially contribute to the emergence of mental health issues during this sensitive period of brain development^8^.

During adolescence, 1 in 5 females initiate hormonal contraceptive (HC) use^9^, primarily in the form of oral contraceptive pills (OCPs), for both contraceptive and therapeutic reasons^10^. HCs contain exogenous hormones, typically ethinyl estradiol and a progestin, which reduce ovarian production of endogenous progesterone^11^ and, to a lesser extent, estradiol^12^. This hormonal reduction leads to the chronic suppression of ovulation. Both of these endogenous hormones are recognized as trophic factors that support brain development and regulate synaptic connectivity^13^. In adults, OCP use has been associated with cortical morphometric brain changes that include lower cortical thickness in regions of the salience and default-mode network (e.g., inferior frontal, lateral orbitofrontal, and cingulate gyri^14^). However, the functional significance of these changes remains unclear, as they were not associated with depressive symptoms^15^. Despite this, research indicates that adolescent HC users might be at an increased risk of developing mental health issues, particularly depressive symptoms, although findings are inconsistent^16^. Notably, the risk for depression appears especially pronounced in young females with attention-deficit/hyperactivity disorder (ADHD) who use HCs^17^, but the underlying mechanism and role of confounding factors (e.g., reason for initiating treatment) remains poorly understood.

Despite the widespread use and extensive testing for safety and efficacy of HCs^18^, their impact on adolescent brain development remains largely unexplored. Leveraging data from the Adolescent Brain Cognitive Development^?^ Study (ABCD Study^®^), we aimed to investigate the effects of hormonal suppression through HC use on brain structure in adolescent females. First, we observed whether hormonal levels of estradiol, testosterone, and dehydroepiandrosterone (DHEA), a precursor to testosterone, were suppressed in HC users (HC+) compared to non-users (HC−). Then, we assessed differences in cortical thickness, surface area, and volume between HC+ and HC− across the brain. Lastly, we investigated associations between hormonal levels and the structural brain measures.

To investigate differences in cortical thickness, surface area, and volume between HC+ and HC−, we leveraged data from 1,234 female adolescents from the ABCD Study from the 4-year follow-up (HC+: *n* = 65; HC−: *n* = 1,169; see **Table 1**). First, using unpaired *t*-tests, we confirmed whether hormonal levels differed between HC+ and HC−. Salivary levels of estradiol, testosterone, and DHEA differed significantly between the two groups. The difference in testosterone was largest (*T* = 4.64, *p* = 0.000004), though all three distributions overlapped considerably (see **Supplementary Figure 1**). Next, a linear mixed-effects model was fitted with group (HC+ vs. HC−) as fixed effect, participant as random effect, and puberty stage, and total intracranial volume (TIV) as covariates. Please see **Supplementary Note 1** for discussion of additional covariates. We observed statistically significant differences between HC+ and HC− in cortical thickness in the bilateral paracentral, superior frontal, medial orbitofrontal, and superior parietal, left precentral, posterior cingulate, lateral occipital, and inferior temporal gyri, as well as the right precuneus. Cortex in all regions was significantly thinner in the HC+ group compared to HC−. After applying False Discovery Rate (FDR) correction^19^ for multiple comparisons, only cortical thickness in the paracentral gyrus remained significantly thinner in HC+ participants than in HC− (left: *p*_FDR_ = 0.0225; right: *p*_FDR_ = 0.0137; see **Supplementary Table 1, Figure 1 Panel A**). HC+ participants exhibited significantly less surface area only in the left postcentral gyrus, but smaller volumes in the bilateral precentral, postcentral, and paracentral gyri, as well as the left lingual gyrus relative to HC− participants. However, results no longer met the threshold of statistical significance after FDR correction. TIV did not differ between the groups.

**Table 1.** Characteristics of individuals included in the analyses.

|  | HC– | HC+ | *p*-value |
| --- | --- | --- | --- |
| **Sample size (*n*)** | 1,169 | 65 |  |
| **Age** | $M = 169.17$ months | $M = 173.31$ months | 0.000087*** |
| **Puberty stage** | $M = 4.18$ | $M = 4.30$ | 0.015* |
| **Combined family income** | $M = 61.65$ | $M = 49.95$ | 0.205 |
| **Area deprivation index** | $M = 39.56$ | $M = 42.79$ | 0.381 |
| **Ethnicity** |  |  |  |
| **White** | 629 (53.81%) | 42 (64.62%) | 0.097 |
| **Black** | 130 (11.12%) | 11 (16.92%) | 0.160 |
| **Hispanic** | 239 (20.44%) | 2 (3.08%) | 0.0001*** |
| **Asian** | 29 (2.48%) | 0 (0%) | n/a |
| **Other** | 142 (12.15%) | 10 (15.38%) | 0.437 |

Note. HC+ = female adolescents using hormonal contraceptives. HC– = female adolescents not using hormonal contraceptives. \* = $p < 0.05$ . \*\*\* = $p < 0.001$

**Figure 1.**
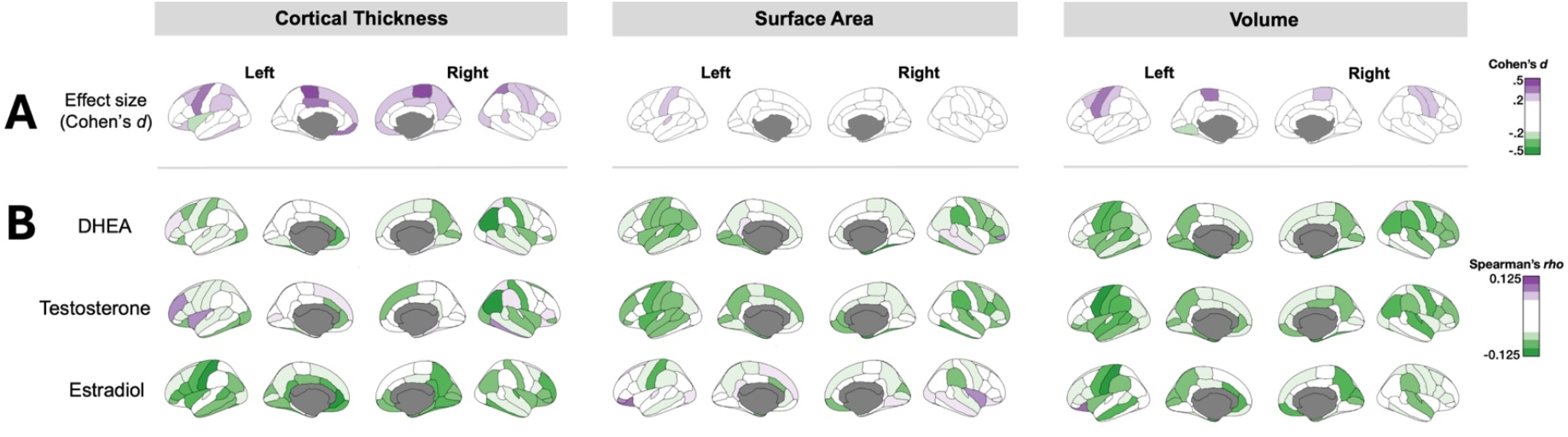
Brain measures of cortical thickness, surface area, and volume. Panel A) Effect sizes (not thresholded for statistical significance) illustrating the impact of hormonal contraceptive (HC) use on cortical thickness, surface area, and volume across the left and right hemispheres. Effect sizes are represented as Cohen’s *d*, with purple indicating lower thickness, surface area, and volume in HC users (HC+) compared to non-users (HC−), green indicating increases in these measurements in HC+. The most pronounced effect is observed as lower cortical thickness bilaterally in HC+, visualized as dark purple in the corresponding regions. Panel B) Partial Spearman correlations of DHEA, testosterone, and estradiol with structural brain measures across both HC users and non-users. Green indicates negative correlations between hormones and brain measures, while purple indicates positive correlations. Correlations were corrected for puberty stage and total intracranial volume (TIV).

Overall, levels of estradiol in both HC+ and HC− youth were negatively correlated with cortical thickness in the right precuneus and left precentral gyrus after adjusting for puberty stage and TIV (see **Supplementary Table 2, Figure 1 Panel B**). Additionally, levels of estradiol, testosterone, and DHEA were negatively correlated with volume in several regions, including the left pre- and postcentral gyri. Surface area of the left postcentral gyrus negatively correlated with levels of estradiol and DHEA. Note that correlations did not remain significant after applying FDR correction. We also evaluated differences in the above-reported correlations between hormone levels and brain measures between HC+ and HC− youth. To test this, separate Spearman correlation coefficients were calculated for each hormone-ROI pair for each group and the slopes of the correlations were compared using Fisher’s *z*-test. However, these comparisons did not survive familywise error correction and were generally small in magnitude. Please see **Supplementary Note 2** for hormone data selection.

In summary, this work contributes to a growing body of literature examining the impact of HCs on brain structure, specifically addressing adolescence as a critical period for neurodevelopment. Using data from over 1,000 youth in the ABCD Study, we revealed hormonal differences and structural differences in HC users compared to non-users, contributing to the understanding of how exogenous hormones might influence the adolescent brain.

Our results demonstrated lower cortical thickness among HC users compared to non-users. Notably, TIV did not differ between HC users and non-users, indicating that observed structural changes were specific to certain brain regions rather than generalized effects of brain size. We observed a statistically significant effect in thickness of the paracentral gyrus, which, in addition to its well-known role in motor function, has recently been linked to a cingulo-opercular network involved in cognitive and physiological control^20^. Likely due to the relatively small number of HC users in this sample (*n* = 65), a number of findings did not survive multiple comparisons corrections, but may still be noteworthy, as they are consistent with previously reported cortical morphometric changes associated with OCP use in adults^14^. Thus, the present work suggests that such findings may extend to an adolescent population, potentially reflecting altered neurodevelopmental trajectories, which emphasizes the importance of studying the effects of hormonal suppression during this critical developmental period.

The observed correlations between estradiol, testosterone, DHEA, and brain structure suggest a subtle but meaningful relationship between hormonal levels and specific brain regions, such as the pre- and postcentral gyri. However, HC use adds complexity to these associations. While we observed significant differences in hormone levels between HC users and non-users—most notably in testosterone—the considerable overlap in hormonal distributions, particularly for estradiol, implies that shifts in hormone levels due to HC use do not necessarily lead to straightforward changes in brain structure. These findings suggest that, although HC use affects hormone levels and potentially brain structure, it does not entirely alter the natural hormone-cortical associations observed in adolescence. Factors such as HC formulation (e.g., progestin type), timing of HC initiation, and individual hormonal sensitivity may modulate these effects, highlighting the need to consider the nuanced ways in which exogenous and endogenous hormones interact to shape brain development. Furthermore, our correlation analyses were sensitive to analytic decisions (such as outlier thresholds), suggesting that hormone-brain associations are complex, while group differences in cortical thickness measurements remained stable across a range of analytic decisions. This emphasizes the need for further research into how individual variations in hormonal sensitivity and HC formulation influence hormone-cortical relationships during adolescence.

Our findings underscore the importance of investigating the neurobiological effects of HCs during the critical period of adolescent neurodevelopment. While HCs have been extensively studied for their contraceptive and therapeutic benefits across all ages, their impact on the brain, especially during development warrants further investigation. We observed structural differences associated with HC use, yet research on the functional significance of these differences is limited. Future work should investigate whether such structural differences reflect adaptive neuroplasticity or compensatory mechanisms in response to hormonal perturbations induced by HCs. Alternatively, these structural differences may signify subtle disruptions in neural circuitry with downstream implications for cognitive processes or emotional regulation. Longitudinal studies incorporating neuroimaging and hormonal assessments could elucidate the trajectory of brain changes associated with HC use over time. Additionally, future research should investigate associations between HC-related brain changes and mental health outcomes in adolescents, particularly regarding mood disorders such as depression^16^. While the underlying mechanisms remain unclear, this work raises questions about the potential synergistic interplay between hormonal modulation, brain structure, and mood regulation. Moreover, comparative analyses with national samples such as from the National Health & Nutrition Examination Survey (NHANES) could provide further context regarding the prevalence and patterns of HC use and their potential implications for population-level brain health.

Several limitations to this study must be noted. First, while this study features a relatively large sample size of 1,234 adolescent individuals, the groups of HC users and non-users were of unequal size (*n* = 65 HC+ versus *n* = 1,169 HC– adolescents, after MRI data quality control). This imbalance may have hindered the detection of true differences between the groups by reducing statistical power and increasing the likelihood of false negatives. Second, assessing HC- and hormone-related differences at the level of the large, anatomical regions of interest (ROIs) defined by the Desikan-Killiany parcellation may limit the detection of endocrine-mediated effects in smaller subregions or functionally delineated brain regions. Given the sample size of HC+ participants, the Desikan-Killiany atlas offers a balanced approach, preserving statistical power. Future studies with larger HC+ cohorts may benefit from voxel-based approaches or more detailed parcellation methods to capture differences in more detail. Third, participants’ reasons for initiating HCs are not known and may confound the observed associations. The HC+ and HC– groups differed demographically (puberty stage, age, ethnicity) and, while we adjusted for these differences, the most robust adjustment would be a longitudinal analysis of the same individuals over time. The ABCD Study is uniquely suited to achieve such a longitudinal analysis, spanning ten years and the entire period of adolescence, as some individuals who are not HC users in the current data wave will undoubtedly initiate HC use in subsequent waves. The cross-sectional design of this current analysis prevents us from drawing causal conclusions about the relationship between HC use and brain structural changes, but future data releases from the ABCD Study, which is both large and longitudinal, will remedy this. Thus, these findings direct the field’s attention toward the important issue of hormone- and contraceptive-related differences in the developing brain and identify it as an area in need of follow-up as more data become available.

In conclusion, our study identifies HC-related differences in childhood cortical morphology and highlights the need for continued research in this area across adolescent development. By elucidating impacts of HC-related hormonal suppression on brain structure and function, or lack thereof, we can better understand the potential risks and benefits associated with HC use during this critical period of brain maturation. Such insights are essential for informing clinical practice and public health policies aimed at optimizing adolescent reproductive, brain, and mental health.

## Online Methods

### Data source

The ABCD Study (https://abcdstudy.org) is a prospective, observational, 10-year longitudinal investigation into brain development, spanning from ages 9 to 10 years at baseline through adulthood and comprising 21 study sites. The study’s rationale and design aspects have been described in detail elsewhere^21^. Approval for the ABCD Study was given from the central Institutional Review Board at the University of California, San Diego and from the local Institutional Review Boards at University of Maryland, Baltimore, University of Colorado, Boulder, University of Minnesota, the Laureate Institute for Brain Research, Oregon Health and Science University, University of Vermont, University of Pittsburgh, Virginia Commonwealth University, University of Rochester, University of Florida, Medical University of South Carolina, University of Michigan, University of Utah, SRI International, University of Wisconsin-Milwaukee, Children’s Hospital of Los Angeles, Florida International University, Washington University in St. Louis, and Yale University. Data used in these analyses (v5.1; https://dx.doi.org/10.15154/z563-zd24) from the 4-year follow-up includes 4,754 children and adolescents enrolled between 2016 and 2022, among whom 1,234 females had both structural MRI and information on HC use. Specifically, the study includes a total of *n* = 1,169 individuals who did not use HCs (HC−) and *n* = 65 individuals who reported current HC use (HC+) and had year 4 follow-up MRI data that passed quality control checks.

### Structural brain imaging

All individuals included in this study underwent 3 Tesla structural MRI scans according to standardized protocols^22^. T1-weighted images were acquired at local study sites and processed at the Data Analysis, Informatics and Resource Center (DAIRC) of the ABCD Study^23^. FreeSurfer v7.1.1 (https://surfer.nmr.mgh.harvard.edu) was used to process the images and obtain brain measures from 34 regions within each brain hemisphere from the Desikan-Killiany atlas^24^.

### Endocrine assessments

Hormone levels of estradiol, testosterone, and DHEA were measured from saliva samples. Analyses were conducted by Salimetrics (https://salimetrics.com). Estradiol was assessed using the Salimetrics^®^ 17β-Estradiol Enzyme Immunoassay Kit (assay range: 1 pg/ml – 32 pg/ml; assay sensitivity 0.1 pg/ml; serum-saliva correlation: 0.80; intra-assay precision, ≤ 8.1%, inter-assay precision: ≤ 8.9%). Testosterone was assessed using the Salimetrics^®^ Testosterone Enzyme Immunoassay Kit (assay range: 6.1 pg/ml – 600 pg/ml; assay sensitivity 1 pg/ml; serum-saliva correlation: 0.96; intra-assay precision, ≤ 6.7%, inter-assay precision: ≤ 13%). DHEA was assessed using the Salimetrics^®^ DHEA Enzyme Immunoassay Kit (assay range: 10.2 pg/ml – 1000 pg/ml; assay sensitivity 5 pg/ml; serum-saliva correlation: 0.86; intra-assay precision, ≤ 5.8%, inter-assay precision: ≤ 8.5%).

### Hormonal contraceptive use

A total of 1,979 adolescents had information available on HC use, measured using a youth survey item which is asked once at each annual visit: *Is your child currently using hormonal birth control (e*.*g. the pill, hormone patch, hormone injection)?* With response choices *“Yes/No”*. We elected to use data from youth rather than parent report because of the possibility that the parent or guardian supervising the study visit may be unaware of contraceptive use.

### Puberty stage and additional covariates

Puberty stage was determined using the Pubertal Development Scale. We used variable name *pds_y_ss_female_category_2* from the latest available wave (Year 4 follow-up). Data used to calculate puberty stages were collected from both parents and youth (variable name: *pds_p_ss_female_category_2, pds_y_ss_female_category_2*, for parent and youth scales, respectively). We noted that the parent and youth scores had only a moderate correlation (*r* = 0.285). Because the staging criteria rely on personal and sensitive information (breast growth and onset of menstruation) we elected to use the youth rather than parent report. Age was also considered as covariate. However, age was collinear with puberty stage, therefore we selected only puberty stage to include in our overall model to avoid multicollinearity (see additional details in **Supplementary Note 1**). Socioeconomic status was also considered as covariate, which was operationalized using Area Deprivation Index (variable name: *reshist_addr1_adi_perc*) and family income (variable name: *demo_comb_income_v2*).

### Statistical approach

Statistical analyses were performed using Python (version 3.9.6) with the libraries *numpy, pandas, scipy*, and *statsmodels*, and visualizations were made using the R package *ggseg*. Demographic variables were compared by Mann-Whitney *U* test due to non-normality in the underlying distributions, except for race/ethnicity, which was compared by Fisher’s Exact Test. In our initial model, effects of HCs on cortical thickness, surface area, and cortical volume were evaluated in a general linear model with group (HC+ vs HC−) as a predictor. Because puberty stage is strongly related to structural brain changes, we tested additional models that included puberty stage and TIV as covariates in the model, which did not change the overall pattern of results. Because the general linear model assumes an underlying Gaussian data distribution, we applied the Shapiro-Wilk test for normality and Breusch-Pagan test for homoscedasticity to the residuals. Some ROIs passed and others failed these assumption checks, therefore we also (1) entered the data into a robust regression, and (2) applied a log transformation to normalize the data, then entered into an ordinary least squares model. The general pattern of results was unchanged regardless of the statistical model employed. Lastly, Spearman correlations and semi-partial Spearman correlations were performed to evaluate the relationship between brain measures and hormone levels, with puberty stage and TIV as covariates. Group differences in hormone levels were evaluated using unpaired *t*-tests. We also evaluated whether the above-reported correlations between hormone levels and brain measures differed between HC+ and HC−. To test this, separate Spearman correlation coefficients were calculated for each hormone-ROI pair for each group and compared using Fisher’s *z*-test. All analyses were FDR-corrected for multiple comparisons^19^. Both corrected and uncorrected results are reported.

All analysis scripts are publicly available on our GitHub repository: https://github.com/HumanBrainZappingatUCLA/ABCDOCPS/

## Acknowledgment

Data used in the preparation of this article were obtained from the Adolescent Brain Cognitive Development (ABCD) Study (https://abcdstudy.org), held in the NIMH Data Archive (NDA). This is a multisite, longitudinal study designed to recruit more than 10,000 children aged 9-10 and follow them over 10 years into early adulthood. The ABCD Study is supported by the National Institutes of Health Grants [U01DA041022, U01DA041028, U01DA041048, U01DA041089, U01DA041106, U01DA041117, U01DA041120, U01DA041134, U01DA041148, U01DA041156, U01DA041174, U24DA041123, U24DA041147]. A full list of supporters is available at https://abcdstudy.org/nih-collaborators. A listing of participating sites and a complete listing of the study investigators can be found at https://abcdstudy.org/principal-investigators.html. ABCD consortium investigators designed and implemented the study and/or provided data but did not necessarily participate in analysis or writing of this report. This manuscript reflects the views of the authors and may not reflect the opinions or views of the NIH or ABCD consortium investigators. The ABCD data repository grows and changes over time. The ABCD data used in this report came from 10.15154/z563-zd24 (Release 5.1). This work was supported by the German Research Foundation (grant number: 5 44183227 to CH), the Northwell Health Advancing Women in Science and Medicine Career Development Award and Educational Achievement Award, as well as the Feinstein Institutes for Medical Research Emerging Scientist Award (to ED), the South-Eastern Norway Regional Health Authority (to CB: 2023037, 2022103).

## References

1. Arain, M. et al. Maturation of the adolescent brain. Neuropsychiatr Dis Treat 9, 449 (2013).

2. Kolb, B. & Gibb, R. Brain Plasticity and Behaviour in the Developing Brain. Journal of the Canadian Academy of Child and Adolescent Psychiatry 20, 265 (2011).

3. Shaw, P. et al. Neurodevelopmental Trajectories of the Human Cerebral Cortex. Journal of Neuroscience 28, 3586–3594 (2008).

4. Gogtay, N. et al. Dynamic mapping of human cortical development during childhood through early adulthood. Proc Natl Acad Sci U S A 101, 8174–8179 (2004).

5. Blakemore, S. J. The social brain in adolescence. Nat Rev Neurosci 9, 267–277 (2008).

6. Herting, M. M., Gautam, P., Spielberg, J. M., Dahl, R. E. & Sowell, E. R. A Longitudinal Study: Changes in Cortical Thickness and Surface Area during Pubertal Maturation. PLoS One 10, (2015).

7. Vijayakumar, N., Op de Macks, Z., Shirtcliff, E. A. & Pfeifer, J. H. Puberty and the human brain: insights into adolescent development. Neurosci Biobehav Rev 92, 417 (2018).

8. Pfeifer, J. H. & Allen, N. B. Puberty initiates cascading relationships between neurodevelopmental, social, and internalizing processes across adolescence. Biol Psychiatry 89, 99 (2021).

9. Abma, J. C. & Martinez, G. M. Sexual Activity and Contraceptive Use Among Teenagers in the United States, 2011–2015. (2011).

10. Armstrong, C. ACOG Guidelines on Noncontraceptive Uses of Hormonal Contraceptives. Am Fam Physician 82, 288–295 (2010).

11. Thorneycroft, I. H. & Stone, S. C. Radioimmunoassay of serum progesterone in women receiving oral contraceptive steroids. Contraception 5, 129–146 (1972).

12. Sullivan, H., Furniss, H., Spona, J. & Elstein, M. Effect of 21-day and 24-day oral contraceptive regimens containing gestodene (60 μg) and ethinyl estradiol (15 μg) on ovarian activity. Fertil Steril 72, 115–120 (1999).

13. McEwen, B. S. & Woolley, C. S. Estradiol and progesterone regulate neuronal structure and synaptic connectivity in adult as well as developing brain. Exp Gerontol 29, 431–436 (1994).

14. Heller, C., Kimmig, A. S., Kubicki, M. R., Derntl, B. & Kikinis, Z. Imaging the human brain on oral contraceptives : A review of structural imaging methods and implications for future research goals. Front Neuroendocrinol 67, 101031 (2022).

15. Petersen, N. et al. Effects of oral contraceptive pills on mood and magnetic resonance imaging measures of prefrontal cortical thickness. Mol Psychiatry 26, 917–926 (2021).

16. Kraft, M. Z. et al. Symptoms of mental disorders and oral contraception use: A systematic review and meta-analysis. Front Neuroendocrinol 72, 101111 (2024).

17. Lundin, C. et al. Hormonal Contraceptive Use and Risk of Depression Among Young Women With Attention-Deficit/Hyperactivity Disorder. J Am Acad Child Adolesc Psychiatry 62, 665– 674 (2023).

18. United Nations. Contraceptive Use by Method 2019. Contraceptive Use by Method 2019 (2019) doi:10.18356/1bd58a10-en.

19. Benjamini, Y. & Hochberg, Y. Controlling the False Discovery Rate: A Practical and Powerful Approach to Multiple Testing. Journal of the Royal Statistical Society. Series B (Methodological) vol. 57 289–300 Preprint at 10.2307/2346101 (1995).

20. Gordon, E. M. et al. A somato-cognitive action network alternates with effector regions in motor cortex. Nature 2023 617:7960 617, 351–359 (2023).

21. Compton, W. M., Dowling, G. J. & Garavan, H. Ensuring the Best Use of Data: The Adolescent Brain Cognitive Development Study. JAMA Pediatr 173, 809–810 (2019).

22. Casey, B. J. et al. The Adolescent Brain Cognitive Development (ABCD) study: Imaging acquisition across 21 sites. Dev Cogn Neurosci 32, 43–54 (2018).

23. Hagler, D. J. et al. Image processing and analysis methods for the Adolescent Brain Cognitive Development Study. Neuroimage 202, (2019).

24. Desikan, R. S. et al. An automated labeling system for subdividing the human cerebral cortex on MRI scans into gyral based regions of interest. Neuroimage 31, 968–980 (2006).

