## Supplemental materials for "Hormonal contraceptive intake during adolescence and cortical brain measures in the ABCD Study"

### Supplementary Material

| | |
|---|---|
| Supplementary Figure 1. | Comparisons of estradiol, testosterone, and DHEA saliva levels in hormonal contraceptive users (HC+) vs non-users (HC−). |
| Supplementary Note 1. | Additional covariate tests and covariate selection procedures. |
| Supplementary Note 2. | Hormone data selection and cleaning methods |
| Supplementary Table 1. | Cortical thickness, surface area, and volume differences in hormonal contraceptive users (HC+) vs non-users (HC−). |
| Supplementary Table 2. | Partial Spearman correlations between brain regions and hormonal levels of estradiol, testosterone, and DHEA. |
| Supplementary Figure 2. | Partial Spearman correlations between hormones and structural brain measures with correction for age (instead of puberty stage) and total intracranial volume (TIV). |

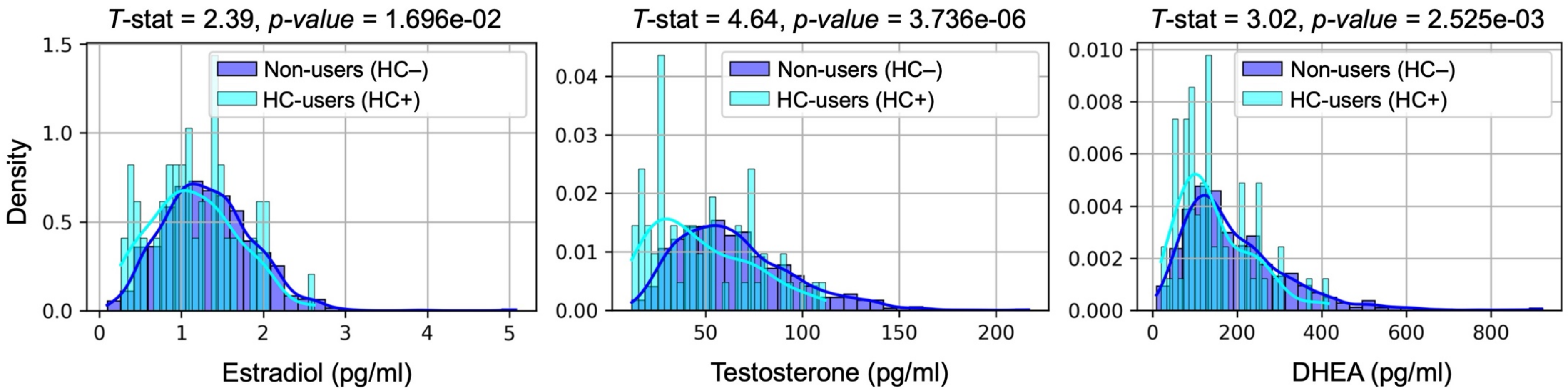


**Supplementary Figure 1. Density histogram of salivary hormone levels in hormonal contraceptives users (HC+) and non-users (HC–).** The figure highlights the central tendency and variability of the data across both groups (HC+ vs HC–). The two groups differed significantly in salivary estradiol, testosterone, and DHEA levels. The difference in testosterone was largest, and all three distributions overlapped considerably. The x-axis represents the hormonal levels, measured in pg/ml. The y-axis represents the density of occurrences within each group, normalized so that the area under the curve equals 1. Histogram bars are overlaid with a smooth density curve.

**Supplementary Note 1. Additional covariate tests and covariate selection procedures.**

In analyses that included puberty stage, we used variable name *pds_y_ss_female_category_2.* Data used to calculate puberty stages were collected from both parents and youth (variable name: *pds_p_ss_female_category_2, pds_y_ss_female_category_2*, for parent and youth scales, respectively). We noted that the parent and youth scores had only a moderate correlation ($r = 0.285$). Because the staging criteria rely on personal and sensitive information (breast growth and onset of menstruation) we elected to use the youth rather than parent report.

We also considered the role of socioeconomic status, which we operationalized using the Area Deprivation Index (variable name: *reshist_addr1_adi_perc*), as in previous investigations[1–3]. We observed a relatively small but statistically significant negative relationship between ADI and cortical thickness that varied by region of interest (*r*s ranging from -0.15456 to 0.07163). However, a Mann-Whitney test (employed due to non-Gaussian data) showed no significant difference in ADI scores between the two study groups ($U$ = 33000.50, $p = 0.382$, [mean HC– = 39.56111, median = 34.0, $n = 1121$; mean HC+ = 42.79365, median = 41.0, $n = 63$), and therefore we ultimately decided to exclude this variable from our models. For completeness, we also applied a Mann-Whitney test to family income (operationalized as variable: *demo_comb_income_v2*), which also did not differ between the two groups ($U = 34506.00$, $p = .205$).

Age also correlated negatively with cortical thickness in most brain regions (range of correlation coefficients: -0.14605 to 0.05566) and differed significantly between the two groups ($U = 27027.0$, $p$ = 8.72462e-05; mean HC– = 169.17 months, mean HC+ = 173.31 months). However, age was also collinear with puberty stage ($r = 0.17979$, $p$ = 2.94915e-10), therefore we selected only puberty stage to include in our primary overall model to avoid multicollinearity. When adding age as a covariate instead, the general pattern of results is similar: the effect of HC on left and right paracentral cortex thickness survives FDR correction.

Some regions (right superior parietal, left superior frontal, left precentral, and left posterior cingulate cortex) remain different only at an uncorrected threshold, as in the puberty-corrected model. In this age-corrected model, right precuneus is replaced by left precuneus, and left insula is also different at the uncorrected threshold.

Given the highly significant difference in age between the two groups and the risk that age could be confounding effects of HCs on cortical thickness, we also took a second approach, age-matched subsampling, to confirm that age differences did not drive the observed effects in the overall model (*i.e.*, testing the effect of HC use on cortical thickness). We trimmed the larger group (HC−) by randomly removing younger participants until the age difference between the two groups was no longer significant ($p = 0.055$). Iterative testing showed that this occurred when 42% of the participants aged 150-170 months were removed from the HC− group.

We then repeated the ordinary least squares analysis with FDR correction to compare cortical thickness between the groups. Despite the substantial reduction in sample size, the effect of HC use on left and right paracentral cortex thickness remained statistically significant. Notably, the $p$-values were slightly smaller after adjusting the age distribution: the $p_{FDR}$ for the left paracentral cortex decreased from 0.0225 to 0.0089, and the $p_{FDR}$ for the right paracentral cortex decreased from 0.0137 to 0.0089. These results provide additional confidence that the observed effect of HC use on paracentral cortical thickness was not an artifact of age differences between the groups.

Finally, ethnicity was somewhat different between the two groups. Fisher's exact test showed that the HC+ group had significantly fewer individuals who identified as Hispanic. Removing this imbalanced group from the analysis altogether (that is, dropping all individuals who identify as Hispanic) did not change the overall results.

***Supplementary References***


1. Vargas, T., Damme, K. S. F. & Mittal, V. A. Neighborhood deprivation, prefrontal morphology and neurocognition in late childhood to early adolescence. *Neuroimage* 220, 117086 (2020).
2. Adise, S., Marshall, A. T., Kan, E., Gonzalez, M. R. & Sowell, E. R. Relating Neighborhood Deprivation to Childhood Obesity in the ABCD Study: Evidence for Theories of Neuroinflammation and Neuronal Stress. *Health Psychology* 42, 868–877 (2022).
3. Ku, B. S. *et al.* Aspects of Area Deprivation Index in Relation to Hippocampal Volume Among Children. *JAMA Netw Open* 7, e2416484–e2416484 (2024).

**Supplementary Note 2: Hormone data selection and cleaning methods**

Hormone data were acquired from table *ph_y_sal_*horm, variable names *hormone_scr_dhea_mean* (DHEA), *hormone_scr_ert* (testosterone), and *hormone_scr_hse* (estradiol). The available hormone data had an unusually high rate of zero values. These could reflect true zeroes (i.e., hormone levels in the participants that were below the limit of detection), or undetectable hormone levels due to errors in sample handling, shipping, processing, analysis, or data management (e.g., erroneously coding missing data as zero). Therefore, based on biologically plausible reference levels, we set a lower boundary for DHEA of 5 pg/mL, for testosterone of 5 pg/mL, and for estradiol of 0 pg/mL. After examining the distribution of data for each hormone, we also removed outliers above 1000, 500, and 1500 pg/mL for each hormone, respectively. Only hormone levels within these boundaries were included in the reported hormone-brain measurement correlations. Notably, these correlations were sensitive to choices in the analytic pipeline, and we present the results of the most conservative analysis.

**Supplementary Table 1.** Cortical thickness, surface area, and volume differences in hormonal contraceptive users (HC+) vs non-users (HC−).

| **Region** | ***p*** | ***p*FDR** |
|---|---|---|
| **Cortical thickness** | | |
| Paracentral (right) | ***0.0002*** | ***0.0137*** |
| Paracentral (left) | ***0.0007*** | ***0.0225*** |
| Superior parietal (right) | ***0.0127*** | n.s. |
| Superior parietal (left) | ***0.0183*** | n.s. |
| Medial orbitofrontal (right) | ***0.0399*** | n.s. |
| Medial orbitofrontal (left) | ***0.0464*** | n.s. |
| Superior frontal (right) | ***0.0481*** | n.s. |
| Superior frontal (left) | ***0.0158*** | n.s. |
| Precuneus (right) | ***0.0222*** | n.s. |
| Precentral (left) | ***0.0143*** | n.s. |
| Posterior cingulate (left) | ***0.0180*** | n.s. |
| Lateral occipital (left) | ***0.0371*** | n.s. |
| Inferior temporal (left) | ***0.0389*** | n.s. |
| **Surface area** | | |
| Postcentral (left) | ***0.0406*** | n.s. |
| **Volume** | | |
| Precentral (right) | ***0.0074*** | n.s. |
| Precentral (left) | ***0.0032*** | n.s. |
| Postcentral (right) | ***0.0408*** | n.s. |
| Postcentral (left) | ***0.0136*** | n.s. |
| Paracentral (right) | ***0.0484*** | n.s. |
| Paracentral (left) | ***0.0078*** | n.s. |
| Lingual (left) | ***0.0221*** | n.s. |

Note. Significant results are indicated in bold and italic. $p_{FDR}$ = *p*-value after correction for multiple comparisons using the False Discovery Rate method. n.s. = not significant.

**Supplementary Table 2.** Partial Spearman correlations between brain regions and hormonal levels of estradiol, testosterone, and DHEA.

| Region | Estradiol | | | Testosterone | | | DHEA | | |
|---|---|---|---|---|---|---|---|---|---|
| | *r* | *p* | $p_{FDR}$ | *r* | *p* | $p_{FDR}$ | *r* | *p* | $p_{FDR}$ |
| **Cortical thickness** | | | | | | | | | |
| Paracentral (right) | -0.024 | 0.479 | n.s. | -0.009 | 0.792 | n.s. | -0.026 | 0.451 | n.s. |
| Paracentral (left) | -0.004 | 0.957 | n.s. | -0.011 | 0.760 | n.s. | 0.002 | 0.960 | n.s. |
| Superior parietal (right) | -0.025 | 0.475 | n.s. | 0.042 | 0.218 | n.s. | 0.034 | 0.320 | n.s. |
| Superior parietal (left) | -0.007 | 0.914 | n.s. | -0.047 | 0.173 | n.s. | -0.033 | 0.341 | n.s. |
| Medial orbitofrontal (right) | -0.055 | 0.108 | n.s. | 0.023 | 0.510 | n.s. | 0.025 | 0.477 | n.s. |
| Medial orbitofrontal (left) | ***-0.083*** | ***0.015*** | n.s. | -0.019 | 0.599 | n.s. | -0.041 | 0.232 | n.s. |
| Superior frontal (right) | -0.025 | 0.475 | n.s. | -0.053 | 0.124 | n.s. | -0.030 | 0.391 | n.s. |
| Superior frontal (left) | -0.007 | 0.850 | n.s. | 0.044 | 0.203 | n.s. | 0.001 | 0.969 | n.s. |
| Precuneus (right) | ***-0.078*** | ***0.024*** | n.s. | -0.023 | 0.510 | n.s. | 0.011 | 0.746 | n.s. |
| Precentral (left) | ***-0.086*** | ***0.012*** | n.s. | -0.044 | 0.205 | n.s. | -0.048 | 0.165 | n.s. |
| Posterior cingulate (left) | -0.064 | 0.062 | n.s. | 0.007 | 0.845 | n.s. | 0.001 | 0.988 | n.s. |
| Lateral occipital (left) | -0.047 | 0.170 | n.s. | -0.055 | 0.108 | n.s. | -0.006 | 0.667 | n.s. |
| Inferior temporal (left) | -0.055 | 0.110 | n.s. | -0.056 | 0.104 | n.s. | -0.055 | 0.107 | n.s. |
| **Surface area** | | | | | | | | | |
| Postcentral (left) | ***-0.082*** | ***0.017*** | n.s. | -0.063 | 0.066 | n.s. | ***-0.070*** | ***0.043*** | n.s. |
| **Volume** | | | | | | | | | |
| Precentral (right) | -0.043 | 0.210 | n.s. | ***-0.088*** | ***0.010*** | n.s. | ***-0.079*** | ***0.021*** | n.s. |
| Precentral (left) | ***-0.081*** | ***0.019*** | n.s. | ***-0.104*** | ***0.002*** | n.s. | ***-0.088*** | ***0.011*** | n.s. |
| Postcentral (right) | -0.036 | 0.295 | n.s. | -0.026 | 0.450 | n.s. | -0.039 | 0.257 | n.s. |
| Postcentral (left) | ***-0.107*** | ***0.002*** | n.s. | ***-0.078*** | ***0.023*** | n.s. | ***-0.092*** | ***0.008*** | n.s. |
| Paracentral (right) | -0.014 | 0.682 | n.s. | -0.030 | 0.380 | n.s. | -0.035 | 0.310 | n.s. |
| Paracentral (left) | -0.049 | 0.157 | n.s. | -0.038 | 0.268 | n.s. | -0.022 | 0.520 | n.s. |
| Lingual (left) | -0.015 | 0.657 | n.s. | -0.043 | 0.217 | n.s. | -0.066 | 0.055 | n.s. |

Note. Spearman correlation corrected for puberty stage and total intracranial volume (TIV). Significant results are indicated in bold and italic. $p_{FDR}$ = *p*-value after correction for multiple comparisons using the False Discovery Rate method. n.s. = not significant.
